# A HEV ORF2 protein-mediated mechanism of hepatitis E-associated kidney disease

**DOI:** 10.1101/2023.09.14.557697

**Authors:** Birgit Helmchen, Anne-Laure Leblond, Daniela Lenggenhager, Jasna Jetzer, Maliki Ankavay, Fritjof Helmchen, Hueseyin Yurtsever, Rossella Parrotta, Marc E. Healy, Amiskwia Pöschel, Enni Markkanen, Nasser Semmo, Martin Ferrié, Laurence Cocquerel, Harald Seeger, Helmut Hopfer, Beat Müllhaupt, Jérôme Gouttenoire, Darius Moradpour, Ariana Gaspert, Achim Weber

## Abstract

Hepatitis E virus (HEV) infection, one of the most common forms of hepatitis worldwide, is often associated with extrahepatic, particularly renal, manifestations. However, the underlying mechanisms are incompletely understood. Here, we report the development of a *de novo* immune complex-mediated glomerulonephritis (GN) in a kidney transplant recipient with chronic hepatitis E. Applying immunostaining, electron microscopy, and mass spectrometry after laser-capture microdissection, we show that GN developed in parallel with increasing glomerular deposition of a noninfectious form of HEV open reading frame 2 (ORF2, capsid) protein secreted in excess. HEV particles or RNA, however, were not detectable. Patients with acute hepatitis E displayed similar but less pronounced deposits. Our results elucidate an immunologic mechanism by which this hepatotropic virus causes variable renal manifestations and establish a link between the HEV ORF2 protein and hepatitis E-associated GN. They directly provide a tool for etiology-based diagnosis of HEV-associated GN as a distinct entity and suggest therapeutic implications.

---

Hepatitis E virus (HEV) infection, one of the most common causes of acute hepatitis, is a major global health problem.^1, 2^ The predominantly enterically transmitted HEV infection has two main epidemiologic patterns that correlate with geographically prevalent HEV genotypes. In resource-limited countries, endemic and epidemic HEV-1 and −2 are transmitted from person to person mainly through contaminated drinking water. In resource-rich countries zoonotic HEV-3 and −4 infections predominate, transmitted mainly through contaminated meat products. Despite its high prevalence in industrialized countries, HEV-3 infection has been underdiagnosed in Europe and North America for many years, in part because of its highly variable clinical presentation.^1-3^ The spectrum ranges from an asymptomatic course to acute, self-limiting hepatitis to acute-on-chronic liver failure in patients with pre-existing liver disease and chronic hepatitis in immunocompromised individuals.^4, 5^

HEV-3 infection in particular has been associated with extrahepatic manifestations, mostly neurological and renal diseases, whose underlying pathomechanisms are still largely unknown.^6^ It is conceivable that, apart from renal injury generally associated with impaired liver function, kidney dysfunction in hepatitis E may be caused - solely or additionally - by HEV- inherent mechanisms. Extrahepatic manifestations develop either directly, i.e. by HEV infection of the respective organs or indirectly, i.e. by immunologic reactions.^6-8^ Histologically confirmed glomerular diseases reported in patients with hepatitis E including membranoproliferative glomerulonephritis (MPGN), with or without cryoglobulinemia, and membranous GN, support an underlying immune-mediated mechanism.^6, 9-11^ However, a direct pathophysiologic link to HEV infection, proving a causal relationship with hepatitis E, has not yet been established.^6^ Central to the understanding of HEV pathogenesis is the genetic organization and life cycle of this positive-strand RNA virus whose genome harbors three main open reading frames (ORF) encoding ORF1 non-structural proteins with viral replicase function, the ORF2 protein, corresponding to the capsid protein and main antigenic structure,^7^ and the ORF3 protein involved in viral particle secretion.^12^ HEV produces different ORF2 isoforms: a non-glycosylated isoform assembled into infectious particles (ORF2i) and glycosylated isoforms (ORF2g/c) secreted in large amounts.^13-15^

Here, we describe the development of *de novo* immune complex-mediated glomerulonephritis (GN) in a patient with chronic hepatitis E, and similar but less pronounced deposits in patients with acute hepatitis E.

Patients’ clinical presentation, histopathologic findings and experimental procedures are detailed in Supplementary Material. Autopsy findings in patient 1 included liver cirrhosis, hepatitis with necrosis, and hepatocytes immunohistochemically positive for the HEV ORF2 protein, confirming hepatitis E. Kidney transplant histology showed persistent proliferative and sclerosing immune complex-mediated GN with a membranoproliferative pattern, consistent with MPGN with immune complexes (IC-MPGN)(Figure 1b), which had been diagnosed in a more subtle form in kidney transplant biopsies taken four and three months before death (Figure 1c). There was no evidence of recurrent IgA nephropathy or antibody-mediated rejection. Remarkably, the renal allograft as well as retrospectively examined previous biopsies with GN showed strong immunohistochemical staining for HEV ORF2 protein, decorating the peripheral capillaries and the mesangium of all glomeruli (Figure 1B, right, and 1C). This indicated virus replication for at least 4 months, thus establishing the diagnosis of chronic hepatitis E.^16^

**Figure 1.**
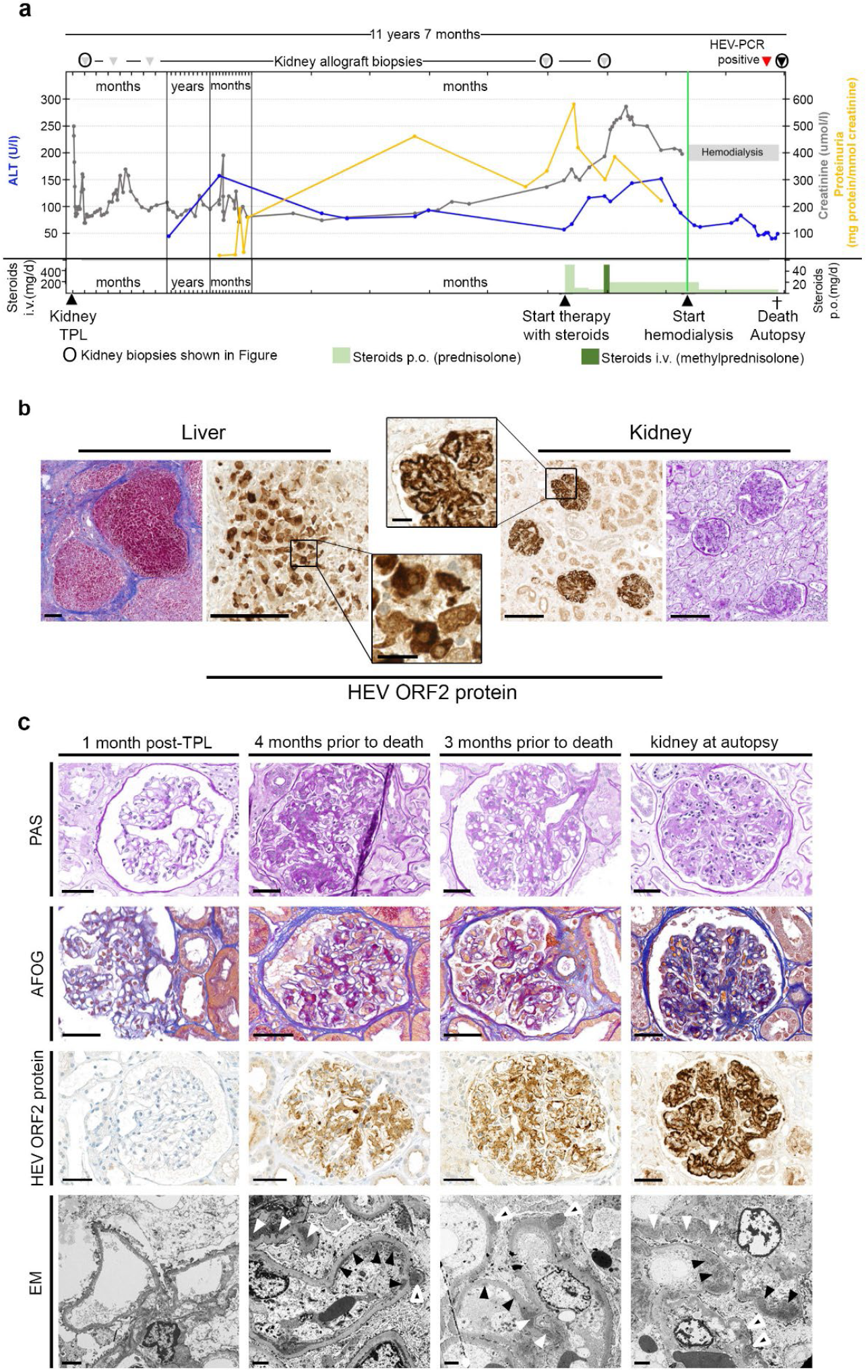
Clinical course, autopsy findings and gradual development of immune complex glomerulonephritis with membranoproliferative pattern in a kidney transplant recipient with hepatitis E. (a) Course of alanine transaminase (ALT, blue), proteinuria (yellow) and creatinine (grey), time points of therapeutic and diagnostic interventions as well as death / autopsy. **(b)** Histology of autopsy liver showing cirrhosis (Masson trichrome stain) and hepatitis with immune reactivity for HEV ORF2 (capsid) protein in hepatocytes (left). Histology of transplant kidney at autopsy showing glomerulonephritis (periodic acid Schiff [PAS] stain) and (extracellular) immune reactivity for HEV ORF2 protein in glomeruli (right). Scale bars in overviews, 200 μm; scale bars in detail, 25 μm. **(c)** Kidney histology. One month post transplantation: inconspicuous glomeruli on light microscopy (PAS and acid fuchsin-Orange G [AFOG] stains), no HEV ORF2 capsid protein deposits, no electron dense deposits on electron microscopy (EM). Four months prior to death (biopsy 4): glomerulus with mild mesangial and endocapillary hypercellularity, segmental sclerosis and prominent podocytes (PAS stain). Mostly mesangial and few glomerular basement membrane protein deposits (AFOG stain). Moderate mesangial and glomerular basement membrane positivity for HEV ORF2 protein. Mesangial (white arrowheads), subendothelial (black arrowheads) and subepithelial (black and white arrowheads) on EM. Three months prior to death (biopsy 5): glomerulus with mild mesangial and endocapillary hypercellularity (PAS stain). Mostly mesangial and few glomerular basement membrane protein deposits (AFOG stain). Moderate to strong mesangial and glomerular basement membrane positivity for HEV ORF2 protein. Mesangial (white arrowheads), subendothelial (black arrowheads) and subepithelial (black and white arrowheads) on EM. Kidney at autopsy: glomerulus with mild mesangial and endocapillary hypercellularity (PAS stain). Mostly mesangial and few glomerular basement membrane protein deposits (AFOG stain). Strong mesangial and glomerular basement membrane positivity for HEV ORF2 protein. Mesangial (white arrowheads), subendothelial (black arrowheads) and subepithelial (black and white arrowheads) on EM. Scale bars in PAS, AFOG, and HEV ORF2 protein images: 50 μm; scale bars in EM images: 2 μm.

HEV RNA was detected by RNA in situ hybridization only in the liver but not in the kidney transplant, as determined in biopsies performed 4 and 3 months before death and at autopsy (Supplementary Figure 1a), indicating that the glomerular HEV ORF2 protein was not associated with HEV virions.

Re-evaluation of the patient’s previous graft biopsies showed a progressive course: Initial subtle mesangial expansion, mild hypercellularity, and immune complex deposition, progressing to a membranoproliferative pattern with endocapillary hypercellularity and significant subendothelial deposits. Reticular aggregates were found in the cytoplasm of endothelial cells at the last graft biopsy. Subendothelial electron dense deposits were confirmed by electron microscopy, which also showed subepithelial and mesangial deposits but no particles suggestive of virions. Prolonged GN was associated with a markedly increased immunohistochemical (IHC) reactivity for the HEV ORF2 protein (Figure 1c), co-localizing with IgG and C3 (Supplementary Figure 1b). Immunofluorescence staining confirmed that colocalization of IgG with HEV ORF2 protein was statistically highly significant (p<10E-10; Figure 2a and Supplementary Figure 2). This was paralleled by worsening renal function and increasing proteinuria (Figure 1a). HEV ORF2 immune complexes were not detected in other organs examined (brain, spleen, heart). Immunoblot analysis using monoclonal antibody (mAb) 1E6 on liver and kidney tissue extracts revealed a band at around 60 kDa, suggesting post-translational modification of the HEV ORF2 protein preserving the epitope region (aa 437-457) (Figure 2b). Combining laser-capture microscopy (LCM) and mass spectrometry (MS) allowed us to further characterize the protein content of the interstitial and glomerular compartments (Supplementary Table 1). LCM/MS analysis of glomeruli revealed, among other fragments, the glomeruli marker podocin and HEV ORF2 protein, notably containing the epitope recognised by mAb 1E6 (Figures 2c-d). In contrast, podocin was not detected in the interstitium, and HEV ORF2 protein only in traces (4 hits versus 241 in the glomerular compartment; Supplementary Table 1). This argued for 1) sufficient differential preparation of glomerular versus interstitial compartments, and 2) principally glomerulus-restricted deposition of HEV ORF2 protein. Furthermore, unlike in HEV-replicating cells and the liver, IHC with mAbs P1H1, P2H1 and P2H2, recognizing only infectious HEV ORF2i^17^ remained negative in the glomeruli, indicating that the glomerular deposits lacked the nonglycosylated, infectious HEV ORF2i (Figure 2e).

**Table 1.**
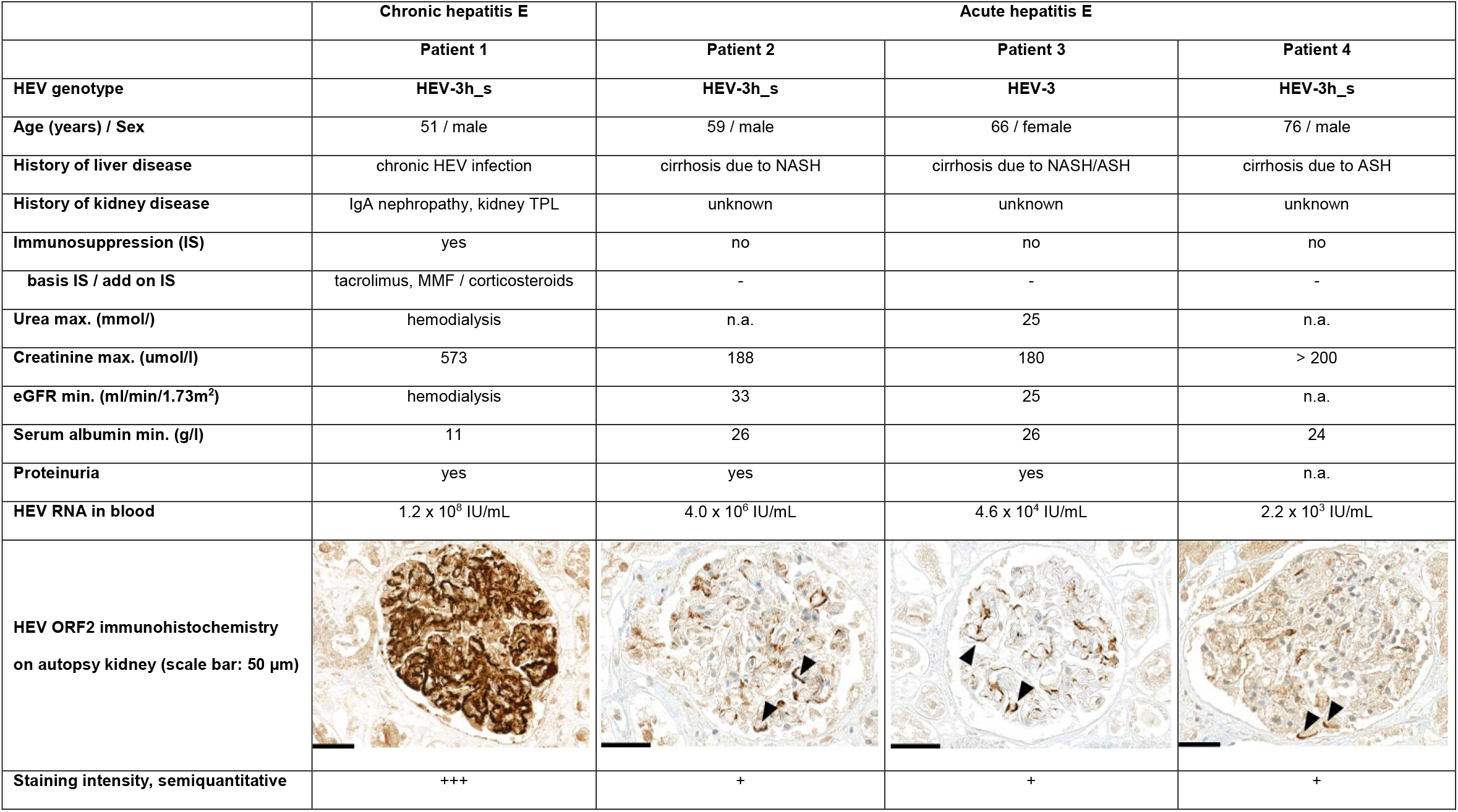
Clinical findings, laboratory values and HEV ORF2 IHC on autopsy kidneys, patients 1-4.^18^. ASH, alcoholic steatohepatitis; HEV, hepatitis E virus; MMF, mycophenolate mofetil; NASH, nonalcoholic steatohepatitis; TPL, transplantation. n.a., information not available

|  | Chronic hepatitis E | Acute hepatitis E |  |  |
| --- | --- | --- | --- | --- |
|  | Patient 1 | Patient 2 | Patient 3 | Patient 4 |
| HEV genotype | HEV-3h_s | HEV-3h_s | HEV-3 | HEV-3h_s |
| Age (years) / Sex | 51 / male | 59 / male | 66 / female | 76 / male |
| History of liver disease | chronic HEV infection | cirrhosis due to NASH | cirrhosis due to NASH/ASH | cirrhosis due to ASH |
| History of kidney disease | IgA nephropathy, kidney TPL | unknown | unknown | unknown |
| Immunosuppression (IS) | yes | no | no | no |
| basis IS / add on IS | tacrolimus, MMF / corticosteroids | - | - | - |
| Urea max. (mmol/l) | hemodialysis | n.a. | 25 | n.a. |
| Creatinine max. (umol/l) | 573 | 188 | 180 | > 200 |
| eGFR min. (ml/min/1.73m<sup>2</sup>) | hemodialysis | 33 | 25 | n.a. |
| Serum albumin min. (g/l) | 11 | 26 | 26 | 24 |
| Proteinuria | yes | yes | yes | n.a. |
| HEV RNA in blood | 1.2 x 10<sup>8</sup> IU/mL | 4.0 x 10<sup>6</sup> IU/mL | 4.6 x 10<sup>4</sup> IU/mL | 2.2 x 10<sup>3</sup> IU/mL |
| HEV ORF2 immunohistochemistry on autopsy kidney (scale bar: 50 µm) |  |  |  |  |
| Staining intensity, semiquantitative | +++ | + | + | + |

**Figure 2.**
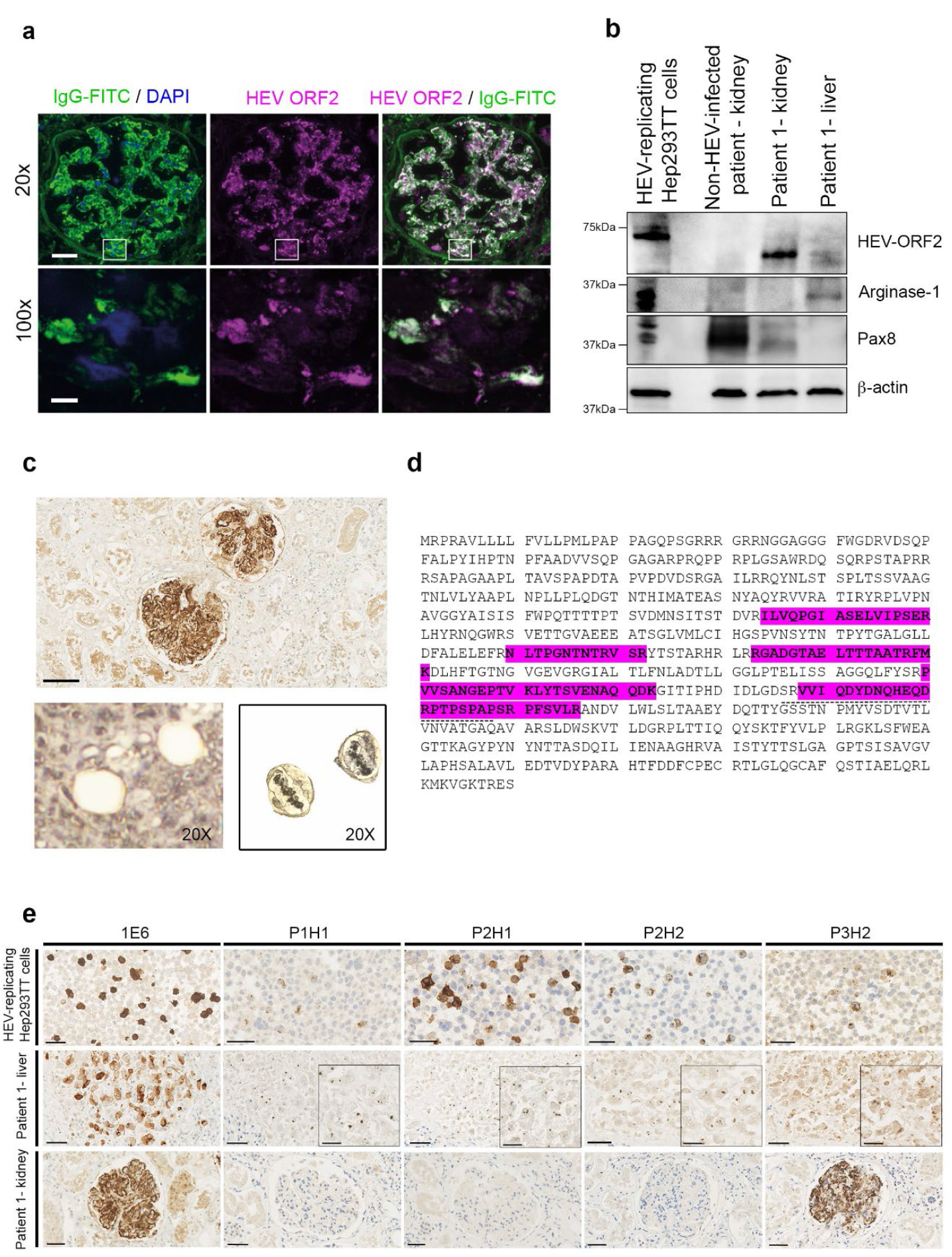
Glomerular deposition of HEV ORF2 protein in a kidney transplant recipient with hepatitis E. (a) Visualization by immunofluorescence staining of a glomerulus from the autopsy transplant kidney (patient 1) with IgG (left: green, FITC stain; DAPI counter-stain, blue) highlighting the co-localization with HEV ORF2 protein (middle: magenta, Alexa546 stain; right: white indicating co-localization). Overview at low magnification (top row, scale bar: 50 µm, 20x) and high resolution images (bottom row, scale bar: 5 µm, 100x) corresponding to the areas indicated by the white boxes. Highly significant IgG/HEV ORF2 protein co-localization was found on the scale of entire glomeruli (20x, Pearson’s correlation coefficient PPC = 0.838 ± 0.039; mean ± s.d., n = 25 glomeruli; p < 10E-10) as well as for small imaging fields (45 – 85 µm side length) acquired at high resolution (100x, Zeiss ApoTome; Pearson’s coefficient 0.668 ± 0.138, n = 16; p < 10E-10). For further glomeruli, please see Supplementary Figure 2. **(b)** Western blot analysis for HEV ORF2 protein, the liver marker Arginase-1, and kidney marker Pax8 in HEV-replicating Hep293TT cells, kidney tissue from a non-HEV-infected patient and patient 1 (Patient 1 - kidney), as well as liver tissue from patient 1 (Patient 1 - liver). **(c)** Laser- capturing of glomeruli positively stained for HEV ORF2 protein by IHC using 1E6 mAb. Scale bar: 100 µm. After excision (lower panel, 20x). Laser-captured glomeruli in the LCM cap (20x). **(d)** Mass spectrometry analysis of the HEV ORF2 sequence derived from laser-captured glomeruli from the transplant kidney tissue of patient 1. Glomerular fragments of HEV ORF2 protein highlighted in magenta. The dashed line depicts the 1E6 epitope. **(e)** Immunohistochemistry using 1E6 and P3H2 mAbs recognizing all the isoforms of HEV ORF2 (ORF2i and ORF2g/c) as well as P1H1, P2H1 and P2H2 mAbs recognizing only infectious HEV ORF2i. HEV-replicating Hep293TT cells, as well as liver (Patient 1 - liver) and kidney tissue (Patient 1 - kidney). Immunoreactivity using P3H2 staining recapitulates IHC results using 1E6 staining in the liver and kidney tissue. Immunoreactivity using P1H1, P2H1 and P2H2 stainings were positive only in Hep293 TT cells expressing ORF2 protein in a replicative context and liver tissue, but not in kidney tissue. Scale bars: 50 μm.

HEV RNA was detected by RT-qPCR in both frozen and FFPE liver specimens. Among tissue specimens from kidney, brain, spleen and heart, HEV RNA was detected only in frozen, but not FFPE specimens (Supplementary Figure 1c). Overall, these findings suggested that the patient had HEV infection of the liver with concomitant viremia and that HEV ORF2 protein aggregates were increasingly deposited in glomeruli. However, they do not provide evidence of a productive HEV infection of kidney cells.

These findings prompted us to examine kidney tissue from additional hepatitis E patients. In three identified cases, all of whom had pre-existing liver cirrhosis and died of acute-on-chronic liver failure in a context of acute hepatitis E,^5, 18^ we found deposits similar as in patient 1, albeit at lower intensities (Table 1). Subtle proliferative glomerular changes with IgG/HEV immune complexes (vizualized by co-immunofluorescence) were detected, consistent with early hepatitis E-associated GN (Supplementary Figure 3).

To determine the significance of the glomerular IgG/HEV ORF2 protein deposits we discovered here for the renal dysfunction observed in association with hepatitis E,^8, 10^ it is important to consider both host and HEV characteristics. Impaired renal function has been reported for HEV-1 and −3, and associated with both acute and chronic hepatitis E,^4, 8^ generally more transient and milder in the acute form.^6^ Its variable presentation includes 1) clinically silent urinary excretion of HEV ORF2 protein with maintained kidney function^19^ 2) transiently impaired kidney function (with or without proteinuria) with resolution following normalization of transaminases,^9, 20^ and 3) subclinical or overt *de novo* immune complex GN with variable outcome including kidney failure.^6, 9^

The development of GN seems to be associated with an impaired immune status.^21^ Accordingly, patient 1 who developed MPGN with bona fide IgG/HEV ORF2 protein deposits was immunocompromized. However, reduced immunocompetence can also be assumed for patients 2-4, who all had liver cirrhosis.^22^ The well-documented temporal association between HEV infection and renal disease, together with the quantitative correlation between ORF2 protein levels and impairment of renal function, argue for a causal relationship between glomerular deposits and renal dysfunction.^9, 19^ In line, we have observed the same type of immune complexes in both acute and chronic hepatitis E, but more pronounced in the latter, in accordance with significantly higher HEV ORF2 protein levels found in sera from chronically as compared to acutely HEV-infected individuals.^23^ In patients 2-4, renal dysfunction was due to hepato-renal syndrome. Glomerular HEV ORF2 protein depositions might have been an aggravating factor. Nevertheless, it is conceivable that in addition to cases of fully developed glomerulonephritis ^6, 9-11^, as in patient 1, glomerular damage associated with more subtle HEV- ORF2 protein deposits, as in patients 2-4, may represent a general early morphological correlate and harbinger of impaired glomerular function in the context of HEV infection.

Based on our observations, we cannot deduce whether HEV ORF2 protein-associated immune-mediated glomerular damage also occurs in acute or subclinical hepatitis E in immunocompetent individuals, in the course of which (transient) impaired renal function has also been described.^2, 8^ However, the host immune status determines the duration of HEV persistence and thus indirectly also the amount of HEV ORF2 protein formed cumulatively.^4, 23^ If this is the decisive factor determining the extent of glomerular damage, it is expected to be lower in acute than in chronic hepatitis E.

HEV antigens can remain detectable in sera of patients with chronic hepatitis E >100 days after clearance of HEV RNA, emphasizing that the presence of HEV ORF2 protein does not necessarily correlate with infectious virions.^23^ It has been shown that HEV exists in urine not only as virions, but also abundantly as free antigen or empty capsid protein, with an obvious discrepancy between the relatively low levels of HEV RNA compared to high levels of HEV ORF2 protein in the urine.^7, 19^ HEV ORF2 protein trapped in the the glomeruli potentially explains the latency observed between viral clearance and restitution of kidney function.^9^

Considering the genetic organization and the HEV life cycle, it is not surprising that the ORF2 (capsid) protein emerges as the key molecule causing extrahepatic manifestations. Unlike ORF1 and ORF3 proteins, the ORF2 protein is produced and secreted in significant excess into the bloodstream, where it exists also in a free form, i.e. not associated with virus particles^13^, remarkably stable and prone to precipitate.^12, 23^ As such, it constitutes the virus’ main immunogenic structure and a potential immunologic decoy.^14, 15^ We noticed that the glomerular HEV ORF2 does not assemble into HEV virions and displays the molecular weight of a truncated HEV ORF2 protein, similar as described in the urine and the stool.^24, 25^ Proteases, as hallmarks of various inflammatory glomerular diseases, may contribute to the cleavage and deglycosylation of the HEV ORF2 protein. Together, these attributes predispose this protein to trigger an immunological reaction.

HEV ORF2 protein IHC and IF, corroborated by Western blotting and MS, allowed us to connect a morphologically variegated and in early stages subtle pattern of glomerular injury (within the spectrum of MPGN) with a specific etiology. Implementation of HEV ORF2 protein immunostaining for routine diagnostics is straightforward^26^ and allows to delineate hepatitis E-associated GN from GN of other causes. This approach is in accordance with the proposed etiology-based classification^27^ and defines hepatitis E-associated GN as a distinct entity.^9^ HEV ORF2 protein immunostaining should be helpful especially in cases in which diagnosis is hampered by limitations of serological testing^1^ and in unsuspected cases of hepatitis E in which extrahepatic manifestations predominate. It may guide therapeutic decisions, especially with regard to immunosuppressive treatment and/or antiviral therapy.^28^

In summary, the discovery of glomerular IgG/HEV ORF2 protein immune complexes may provide a mechanistic explanation for how this antigen triggers renal disease, in particular immune complex GN. Our findings potentially explain several still incompletely understood observations on hepatitis E-associated renal disease, and establish a molecular link between HEV infection and kidney dysfunction.^6^ Finally, they propose ORF2 protein immunostaining as a diagnostic tool for hepatitis E-associated GN, especially in IC-MPGN of clinically unclear origin.

## Supporting information

Supplementary Appendix with figures and tables

## Acknowledgments

We would like to thank Angela Broggini, PhD, Rita Bopp, André Fritsche, Carmen Gavrisan, Marcel Glönkler, Doris Guntersweiler and Michael Reinehr, MD for their excellent assistance by performing the autopsy, immune stains and electron microscopy analyses and providing reagents, respectively.

## List of Abbreviations

HEV,: hepatitis E virus;
GN,: glomerulonephritis;
gt,: genotype;
MPGN,: membranoproliferative glomerulonephritis;
ORF,: open reading frame;
mAb,: monoclonal antibody;
IHC,: immunohistochemistry;
PEN,: polyethylene naphthalate;
LCM,: laser-capture microscopy;
MS,: mass spectrometry.

