## Supplementary Appendix with figures and tables for "A HEV ORF2 protein-mediated mechanism of hepatitis E-associated kidney disease"

**Running title**: Hepatitis E-associated kidney disease

#### TABLE OF CONTENTS

SUPPLEMENTARY FIGURES
SUPPLEMENTARY TABLES
DETAILED CLINICAL INFORMATION, PATIENTS 1-4
  Patients and tissue samples
  Patient 1
  Patient 2
  Patient 3
  Patient 4
DETAILED HISTOPATHOLOGIC DESCRIPTION, PATIENTS 1-4
  Patient 1 – kidney specimens
  Patient 1 – liver specimen
  Patients 2-4 – kidney specimens
  Patients 2-4 – liver specimens
DETAILED DESCRIPTION OF THE METHODS
  Histopathologic evaluation of tissue samples
  Immunohistochemistry (IHC)
  Immunofluorescence
  Transmission electron microscopy (EM)
  In situ hybridization (ISH) for HEV RNA
  Laser capture microscopy (LCM)
  Laser-captured sample preparation for mass spectrometry (MS)-based protein identification
  Image processing
  Western blotting
  Molecular testing: qRT-PCR for HEV
  Molecular testing: HEV genotyping
  Serological assay for anti-HEV IgG and IgM
REFERENCES

### SUPPLEMENTARY FIGURES

#### Supplementary Figure 1

**a**

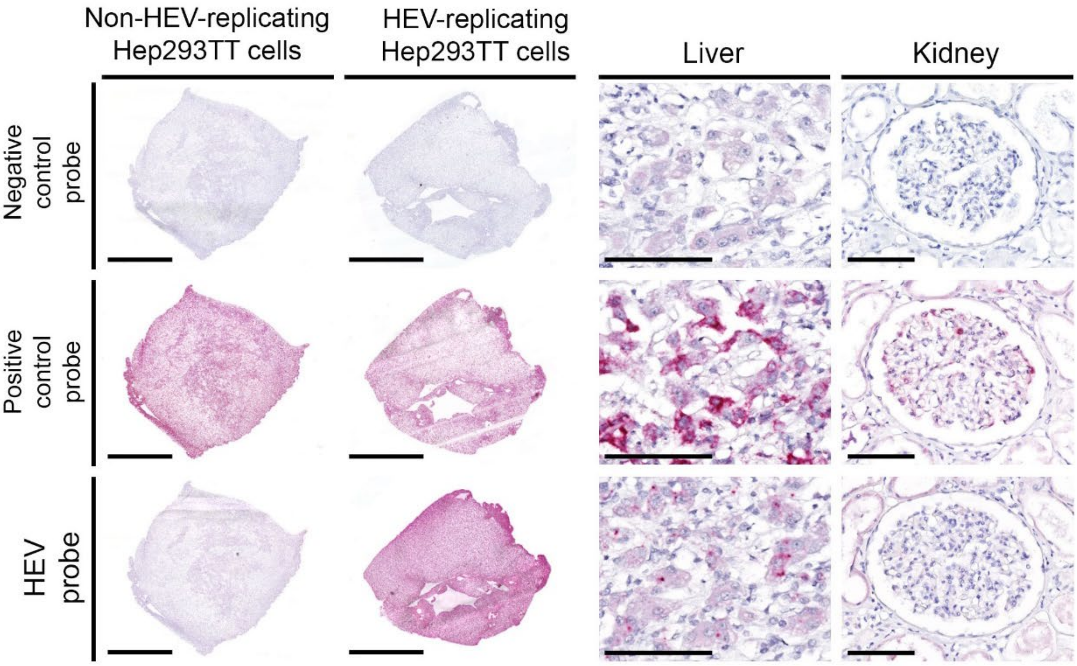

**b**

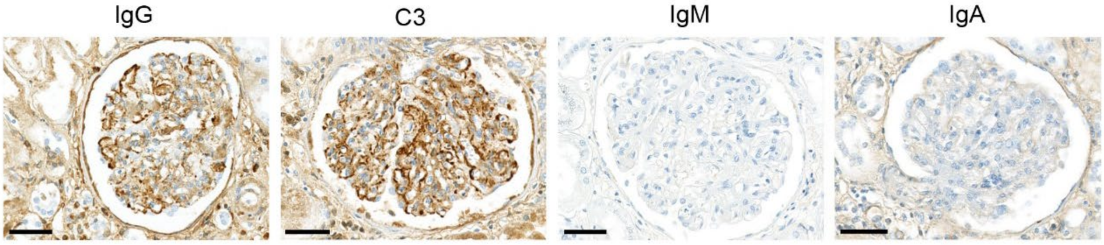

**c**

| Organ | Specimen | Material | HEV ORF2 Ct value | ACTB Ct Value | Result |
|---|---|---|---|---|---|
| **Liver** | Autopsy | FF | 25.26 | 28.88 | Positive |
| | Autopsy | FFPE | 27.73 | 32.67 | Positive |
| **Kidney** | Autopsy | FF | 34.22 | 23.91 | Positive |
| | Autopsy | FFPE | n.d. | 29.63 | Negative |
| | Biopsy 1 | FFPE | n.d. | 28.52 | Negative |
| | Biopsy 4 | FFPE | n.d. | 28.77 | Negative |
| | Biopsy 5 | FFPE | n.d. | 27.5 | Negative |
| **Heart** | Autopsy | FF | 32.76 | 26.29 | Positive |
| **Spleen** | Autopsy | FF | 32.21 | 23.96 | Positive |
| **Brain** | Autopsy | FF | 29.44 | 22.33 | Positive |
| | Autopsy | FFPE | n.d. | 29.18 | Negative |

FFPE: Formalin-Fixed Pararffin-Embedded, FF: Fresh Frozen
Biopsy 1, 1 month P.T.L; Biopsy 4, 4 months prior to death; Biopsy 5, 3 months prior to death; n.d., not detectable

**Supplementary Figure 1. Further autopsy findings, patient 1 (a)** In situ hybridization (ISH) using negative control (upper panel), positive control (middle panel) and HEV-specific (lower panel) probes, on non-HEV-replicating Hep293TT cells and HEV-replicating Hep293TT cells[1] as well as autopsy liver and kidney tissue (scale bars: 2 mm; liver: 100 µm; kidney: 100 µm). **(b)** Immunohistochemistry for IgG with moderate (2+) mesangial and glomerular basement membrane deposits, C3 with moderate (2+) mesangial and glomerular basement membrane deposits, IgM and IgA with negative staining patterns (scale bars: 50 µm). **(c)** qRT-PCR testing for HEV ORF2 and Actin B (ACTB) RNA. Ct value: per 40 ng cDNA input; n.d. not determined.

Supplementary Figure 2

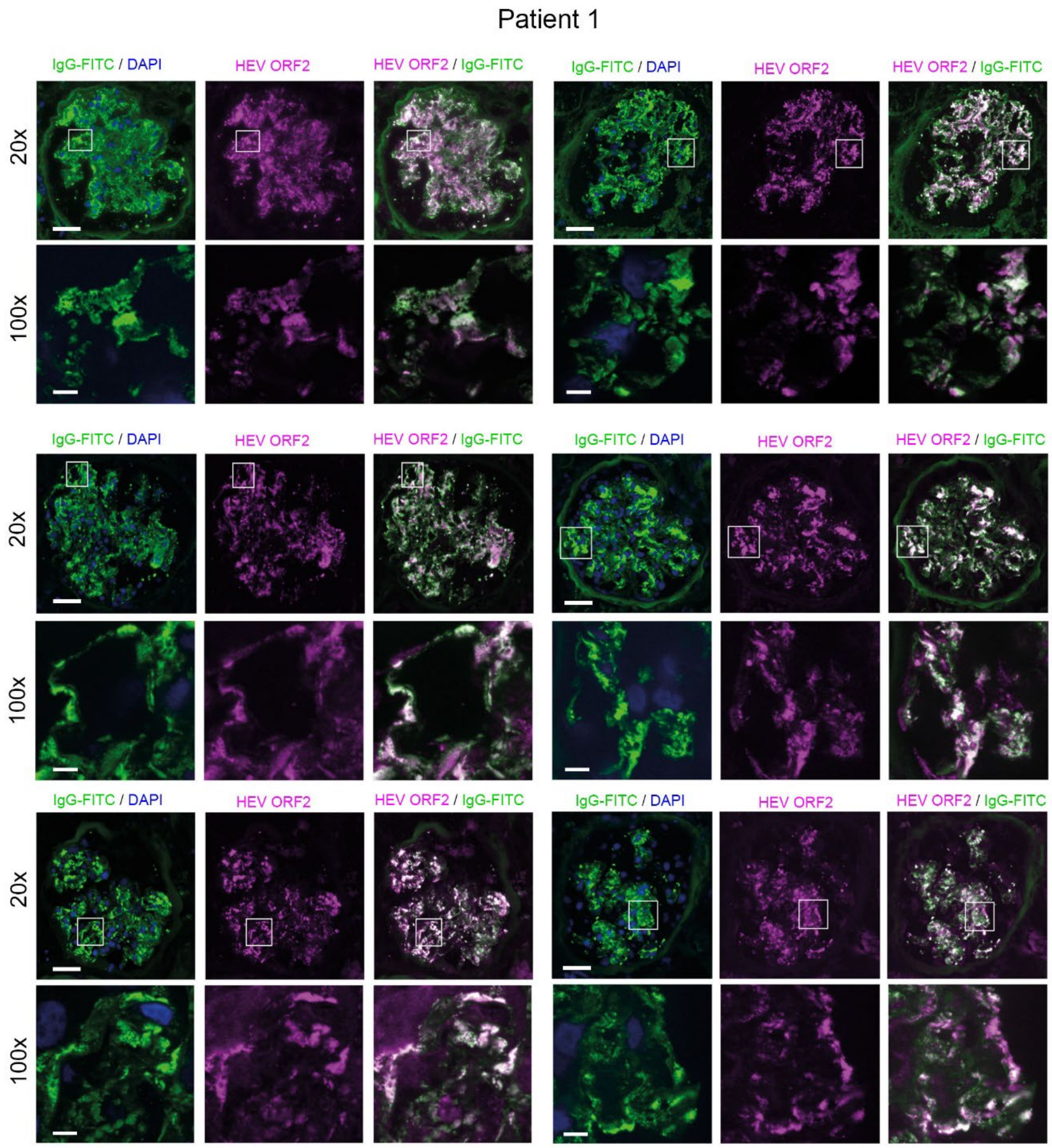

**Supplementary Figure 2: Glomerular IgG/HEV ORF2 co-localization, patient 1**

Visualization by immunofluorescence staining of six glomeruli from the autopsy transplant kidney (patient 1) as in Figure 2A. IgG (left: green, FITC stain; DAPI counter-stain, blue) highlighting the co-localization with HEV ORF2 (middle: magenta, Alexa546 stain; right: overlay with white indicating co-localization). For each glomerulus, overview at low magnification (top rows, scale bar: 50 µm, 20x) and high-resolution images (bottom rows, scale bar: 5µm, 100x) corresponding to the areas indicated by the white boxes.

Supplementary Figure 3

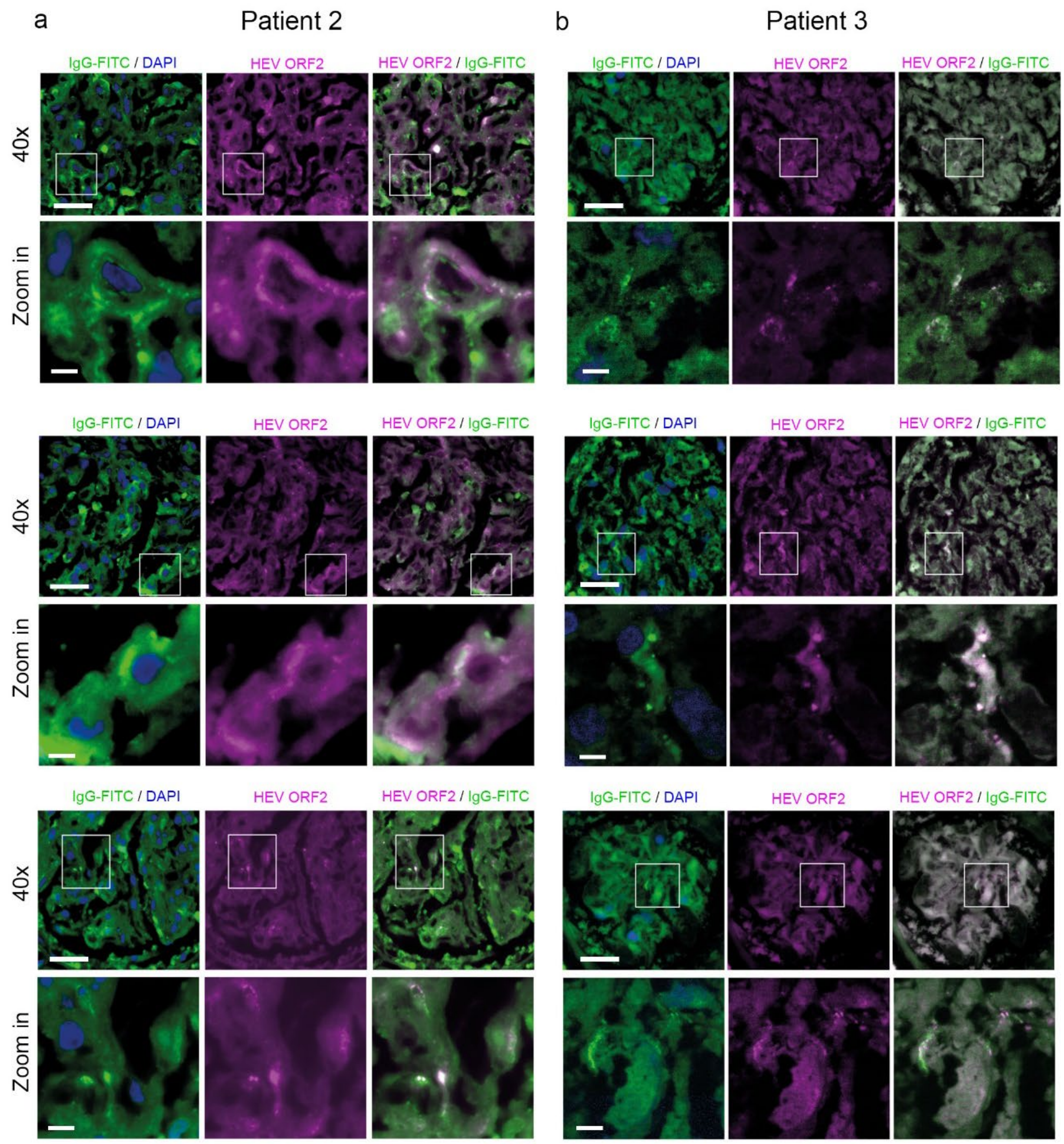

**Supplementary Figure 3: Glomerular IgG/HEV ORF2 co-localization, patients 2 and 3**

**(a)** Visualization by immunofluorescence staining of three glomeruli from patient 2, for which sparse, partially co-localized staining was found. IgG (left: green, FITC stain; DAPI counter-stain, blue) and HEV ORF2 (middle: magenta, Alexa546 stain; right: overlay with white indicating co-localization). For each glomerulus, overview at low magnification (top rows, scale bar: 30 µm, 40x) and zoom-in images (bottom rows, scale bar: 5 µm, zoom in) corresponding to the areas indicated by the white boxes. **(b)** Same as in A for three glomeruli from patient 3. Here, high-resolution images (bottom rows) were obtained with the 100x objective and the ApoTome.

### SUPPLEMENTARY TABLES

| Patient 1 - kidney | | Biopsy 1 | Biopsy 2 | Biopsy 3 | Biopsy 4 | Biopsy 5 | Autopsy |
|---|---|---|---|---|---|---|---|
| | | 1 month post-TPL | 3.5 months post-TPL | 6.5 months post-TPL | 11 years and 3 months post-TPL | 11 years and 4 months post-TPL | 11 years and 7 months post-TPL |
| **Source** | | University Basel Hospital, Switzerland | | | Cantonal Hospital Aarau, Switzerland | | Zürich University Hospital, Switzerland |
| **Material** | | FFPE<br>FF | | | FFPE<br>FF<br>Glutaraldehyde fixed | | |
| **Histological description** | | **Refer to supplementary materials, pages 6-11** | | | | | |
| **According to original reports** | **IHC** | (-) SV40 | (+) SV40 | | Not mentioned | | Not applicable |
| | **IF** | (-) IgA, IgM, C3<br>(-) C4d in peritubular capillaries | (-) IgG, IgA, IgM, and C3<br>(-) C4d in peritubular capillaries | (-) IgA, IgM, IgG,<br>(-) C4d in peritubular capillaries | (-) IgA, IgM, kappa and lambda, C4d<br><br>(+) IgG and C3, granular positivity in GBM and mesangium | (-) C4d, IgM<br>(+) glomerular IgG, kappa and lambda<br><br>IgA and C3 not evaluable | Not applicable |
| | **Diagnosis** | Mild focal tubular atrophy and interstitial fibrosis | Polyomavirus nephropathy | Polyomavirus nephropathy | Proliferative and focal sclerosing immune complex-mediated GN | | Not applicable |
| **USZ workup** | **IHC** | (-) SV40 | (+) SV40 | (-) SV40* | (-) IgA and IgM, SV40<br>(+) IgG and C3 in GBM and mesangium | (-) IgA, C4d<br>(+) IgG, C3 in GBM and mesiangium | (-) SV40, IgA, IgM<br>(+) IgG, C3 in GBM and mesangium |
| | **HEV ORF2 IHC** | (-) HEV ORF2 (Figure 1c) | | | (+) HEV ORF2 (Figure 1c) | | |
| | **IF** | Frozen tissue not available | | | | | (-) IgA, C4d<br>(+) IgG, kappa and lambda, C3 |
| | **EM** | No electron dense deposit | Not done | | Electron dense inhomogenously osmiophilic mesangial, subendothelial and subepithelial deposits. No substructure. | | |
| | **Diagnosis** | Mild focal tubular atrophy and interstitial fibrosis. | Polyomavirus nephropathy. | Polyomavirus nephropathy. Mild arteriolosclerosis. | Proliferative and sclerosing immune complex-mediated GN consistent with HEV-associated GN | | Proliferative and sclerosing immune complex-mediated GN with a membranoproliferative pattern, consistent with HEV-associated membranoproliferative GN with immune complexes |
| | | No evidence for antibody-mediated rejection or recurrent IgA nephropathy | | | | | |

**Supplementary Table 1**: Summary of initial and newly performed analysis and subsequent diagnostics for biopsies and autopsy materials of patient 1.

Post-TPL, post-transplantation, USZ, University Hospital Zurich, FFPE: formalin-fixed, paraffin-embedded, FF: fresh frozen, GBM: glomerular basement membrane, EM: electron microscopy, GN: glomerulonephritis

*HE morphology was indicative of BK infection, suggestive for resolving BK infection.

| | **Non-HEV-infected patient** | | **Patient 1** | |
|---|---|---|---|---|
| **Total spectrum count** | **Interstitium** | **Glomeruli** | **Interstitium** | **Glomeruli** |
| HEV ORF2 | 0 | 0 | 4 | **241 ± 47** |
| Podocin | 0 | **17 ± 2** | 0 | **13 ± 4** |
| Cytokeratin 7 | 33 | 28 | 20 | 29 ± 1 |
| Complement 3 | 74 | 114 ± 2 | 23 | 236 ± 46 |
| Collagen 1A1 | 12 | 22 ± 1 | 27 | 36 ± 2 |

**Supplementary Table 2:** Mass spectrometry results of laser-captured interstitium and glomeruli from a non-HEV-infected patient and patient 1. Protein extract of laser-captured glomeruli from patient 1 contains HEV ORF2 capsid protein, corroborating the IHC results. Protein extract of all laser-captured glomeruli was enriched in glomerular marker podocin[2], that was absent in the laser-captured neighboring interstitial tissue, indicating a sufficient separation of the renal compartments using laser-capture microscopy. The renal markers cytokeratin 7, complement 3 and collagen 1A1 were detectable in both compartments.[3] Results are expressed as total spectrum count with over 95% probability in mean ± s.d.

### DETAILED CLINICAL INFORMATION, PATIENTS 1-4

#### Patients and tissue samples

This study was approved by the internal review board of the University Hospital Zurich and the Cantonal Ethics Committee of Zurich, Switzerland (KEK-ZH-Nr. 2013-0504).

#### Patient 1

This male patient underwent renal transplantation at the age of 40 years for end-stage renal disease due to rapidly progressive crescentic IgA nephritis. Initial immunosuppression consisted of tacrolimus, mycophenolate mofetil (MMF) and sirolimus. Declining renal function prompted allograft biopsy four weeks post transplantation with focal tubulointerstitial fibrosis but no evidence of rejection or calcineurin inhibitor toxicity. His subsequent course was complicated by BK virus (BKV) nephropathy noted in a protocol biopsy 3.5 months post-transplant (post-TPL) and in a protocol biopsy 6.5 months post-TPL. Immunosuppression was reduced. Following BKV clearance and stabilization of renal function baseline, immunosuppression consisted of tacrolimus and MMF. Two years post transplantation the patient developed type 2 diabetes mellitus and was treated with oral antidiabetic agents.

Ten years after transplantation deranged liver function tests were noted (AST: 135 U/l, ALT: 156 U/L, 19 months prior to death). Hepatitis B surface (HBs) antigen and anti-hepatitis C virus were negative, anti-HBs and anti-hepatitis B core positive, consistent with previously known resolved hepatitis B. Drug history and history of alcohol consumption were inconspicuous. Further diagnostic work-up revealed liver cirrhosis of at the time unknown etiology, with splenomegaly, esophageal varices and ascites.

Eleven years and 3 months post-TPL (i.e. 4 months prior to death) rising creatinine levels, proteinuria and microhematuria triggered a renal transplant biopsy to rule out recurrent IgA nephropathy. A diagnosis of *de novo* immune complex glomerulonephritis (GN) positive for IgG and C3 on immunofluorescence was made and confirmed by the presence of subendothelial, mesangial and subepithelial electron dense deposits on electron microscopy. A focal interstitial inflammatory infiltrate and tubulitis were interpreted as being associated with the glomerulonephritis, although borderline changes suspicious for acute T-cell mediated rejection could not be excluded. When the patient was hospitalized for 8 days to adjust his immunosuppressive therapy, he presented with exertional dyspnea, night sweats and weight loss of 16 kg over the last six months upon admission. Steroid therapy was initiated. Given the resolved hepatitis B virus infection, entecavir prophylaxis was initiated. Two weeks later the patient was readmitted for progressive renal failure and persistent nephritic syndrome. Repeat renal biopsy (3 months prior to death, 11 years and 4 month post-TPL) revealed persistent immune complex GN without evidence of rejection. Complement components C3/C4 were within normal limits. The patient was treated with intravenous (IV) pulses followed by oral steroids. Four weeks later he was admitted for renal allograft failure. Thrice-weekly hemodialysis was initiated. Transjugular liver biopsy was reported as chronic hepatitis with mild inflammatory activity on a background of bridging fibrosis or cirrhosis.

Two weeks before his death the patient was readmitted for further evaluation of deteriorating liver function. He had noted increased jaundice and pruritus. He also reported dyspnea and productive cough. Intravenous albumin for severe hypoalbuminemia and vitamin K substitution were initiated. Serologic tests were positive for anti-HEV IgG and negative for anti-HEV IgM (VIDAS® ANTI-HEV IgG and IgM assays, BioMérieux, France). Eight days before death the patient was transferred to a tertiary care center for evaluation of liver transplantation. The following tests were negative / inconspicuous: α1-antitrypsin, anti-double stranded DNA antibodies, rheumatoid factor, ANCA, MPO-ANCA, PR3-ANCA, anti-glomerular basement membrane antibodies, hepatitis A IgM antibodies, hepatitis B surface antigen, hepatitis B core IgM antibody, hepatitis B e-antigen antibodies, hepatitis D virus antibodies, hepatitis C virus antibodies, HIV-1 and HIV-2 antigen and antibody screen, human T-cell lymphotropic virus types I and II antibody, PCR for HBV DNA, HCV RNA, CMV DNA and BKV DNA. Tests for hepatitis A IgG antibodies, hepatitis B surface antibodies, hepatitis B core antibodies, hepatitis E virus IgG antibodies were positive. In contrast to previous testing, also hepatitis E virus IgM antibodies were now detectable (Enzyme ImmunoAssays provided by Dia. Pro Diagnostic Bioprobes Srl, Italy) probably due to the different serological tests used.[4] Over the next days, the patient became hemodynamically unstable and developed hepatic encephalopathy. A chest CT scan showed bilateral infiltrates consistent with nosocomial versus aspiration pneumonia. Despite maximal intensive care treatment, he developed multi-organ dysfunction syndrome. In the absence of therapeutic options, supportive care was initiated. The patient died the following day, 11 years and 7 months after kidney transplantation. The HEV RNA PCR result, which became available only after the patient's death, was positive with very high viremia of 1.2 x $10^8$ IU/mL.

#### Patient 2

This 59-year-old male patient with known liver cirrhosis due to nonalcoholic steatohepatitis, metabolic syndrome and history of coronary artery disease, was admitted to a regional hospital because of acute-on-chronic liver disease. Initial laboratory tests revealed elevated serum aminotransferases and cholestatic parameters, hyperbilirubinemia, decreased albumin and fibrinogen levels as well as prolonged prothrombin time. Further analysis revealed positive anti-HEV IgG and IgM antibodies and HEV RNA of 4.0 x $10^6$ IU/mL, consistent with acute hepatitis E. Transient deterioration of renal function (maximum creatinine level 150 µmol/L, minimal GFR 42 mL/min/1.73 $m^2$) was interpreted as stage1 acute kidney injury. Renal function improved after treatment with IV albumin. After initial improvement of hepatic and renal parameters, the clinical course was complicated by an upper gastrointestinal bleeding, which led to an acute deterioration of hepatic and renal function (max. creatinine 188 µmol/L, minimal eGFR 33 mL/min/1.73 $m^2$). The patient developed hepatic encephalopathy and was transferred to a tertiary care center. At admission, the patient was anuric and required hemodialysis which was attributed to hepato-renal syndrome. A transjugular liver biopsy confirmed the clinically suspected cirrhosis. Three days after admission, the patient experienced severe hemodynamic instability due to recurrent bleeding from duodenal ulcerations. Four days after admission to the tertiary care center, the patient developed severe distributive and septic shock with marked coagulopathy and died. Autopsy confirmed acute-on-chronic liver disease and duodenal ulcerations.

#### Patient 3

This 66-year-old female patient with a history of type 2 diabetes mellitus (treated with oral antidiabetic agents) was admitted to a Cantonal Hospital for suspected acute-on-chronic liver disease. A CT scan revealed liver cirrhosis with mild splenomegaly and moderate ascites in all four quadrants. Laboratory tests revealed elevated levels of serum aminotransferases, cholestatic parameters, hyperbilirubinemia, increased INR and elevated ammonia levels. The hemogram was normal, including a normal platelet count. There were no evidence of hepato-renal syndrome with only slightly elevated urea (9.9 mmol/L), but otherwise normal renal retention parameters (creatinine 70 µmol/L, eGFR 95 mL/min/1.73 $m^2$). Hepatopathy screening was positive for HEV RNA (4.6 x $10^4$ IU/mL) as well as HEV IgG and IgM antibodies, consistent with acute hepatitis E on top of pre-existing liver disease. Antiviral therapy with ribavirin was initiated. A transjugular liver biopsy confirmed cirrhosis. During hospitalization, the patients renal function deteriorated considerably from day 9 after admission, which was interpreted as hepato-renal syndrome (values two weeks after admission: creatinine 180 µmol/L, eGFR 25 mL/min/1.73 $m^2$, urea 25 mmol/L). The patient subsequently developed hepatic encephalopathy and was transferred to a tertiary care center. Within hours after admission, rapid neurological and respiratory deteriorations were noted. After anuric kidney failure, progressive encephalopathy and respiratory failure with pulmonary edema, the patient experienced severe distributive shock with marked coagulopathy and succumbed one day later. Autopsy confirmed acute-on-chronic liver disease.

#### Patient 4

This 76-year-old male patient was admitted to a tertiary care center for suspected acute-on-chronic liver disease. He had a history of alcoholic liver cirrhosis, metabolic syndrome with obesity, insulin resistance and arterial hypertension. Laboratory tests revealed elevated serum aminotransferases and cholestatic parameters, hyperbilirubinemia and hypoalbuminemia (24 g/L). A transjugular liver biopsy confirmed liver cirrhosis. Further laboratory tests revealed positive anti-HEV IgG and IgM antibodies as well as HEV RNA of 2.2 x $10^3$ IU/mL, consistent with acute hepatitis E. Antiviral therapy with ribavirin was initiated. Ribavirin treatment was discontinued five days later due to worsening renal function. From day 9 after admission hepatic encephalopathy as well as the renal function worsened, the latter diagnosed as hepato-renal syndrome. From day 11 after admission, the renal retention parameters worsened significantly (creatinine > 200 µmol/L), prompting therapy with terlipressin and albumin. However, there was no improvement, requiring initiation of hemodialysis. In view of the dramatic deterioration, treatment was switched to comfort care, and the patient died 16 days after admission to the tertiary care center. An autopsy was performed which confirmed the acute-on-chronic liver disease.

Clinical findings obtained in patients 2-4 with respect to their liver phenotype were recently described in Vieira Barbosa et al.[5]

### DETAILED HISTOPATHOLOGIC DESCRIPTION, PATIENTS 1-4

#### Patient 1 – kidney specimens

Staining, electron microscopy analysis and diagnostics are summarized in supplementary table 1.

**Allograft kidney biopsy (1 month post-transplantation (post-TPL), Figure 1c):**

Light microscopy revealed a core of renal cortex containing up 12 glomeruli all with unremarkable features on light microscopy. Minimal interstitial fibrosis and tubular atrophy comprising <5% of the cortex, few assessable arterioles and arteries with unremarkable features were found. No peritubular capillaritis was found.

Immunofluorescence (as stated in the original biopsy report) revealed negative staining results for IgA, IgM and C3 in the glomeruli. IgG was not evaluable. C4d was negative in peritubular capillaries.

Immunohistochemistry (as stated in the original biopsy report): SV40 was negative.

Immunohistochemistry (performed for this study): HEV ORF2: Negative.

Electron microscopy was not performed for initial evaluation.

Electron microscopy (performed for this study) on renal tissue obtained from the paraffin block revealed no electron dense deposits.

**Diagnosis: Mild focal tubular atrophy and interstitial fibrosis. No evidence of rejection, polyomavirus nephropathy or recurrent IgA nephropathy.**

**Allograft kidney biopsy (3.5 months post-TPL):**

Light microscopy: Renal cortex with 18 glomeruli, 2 hyalinized. Focal interstitial edema and predominantly lymphocytic infiltrates comprising approximately 10-20% of the cortex, with tubulitis and intranuclear inclusions in the tubular epithelium typical for polyomavirus. Focal interstitial fibrosis and tubular atrophy, involving approximately 5-10% of the cortex. One artery with mild intimal fibrosis and elastosis. Some arteries and arterioles with swollen myocytes with pale cytoplasm.

Immunofluorescence (as stated in the biopsy report): IgA, IgG, IgM and C3 negative in glomeruli. C4d negative in peritubular capillaries.

Immunohistochemistry (as stated in the original biopsy report): SV40 focally positive in nuclei of tubular epithelial cells.

Immunohistochemistry (performed for this study): HEV ORF2: Negative.

Electron microscopy was not performed.

**Diagnosis: Polyomavirus nephropathy. Minimal interstitial fibrosis and tubular atrophy. Minimal signs of calcineurin-inhibitor-associated toxicity on praeglomerular vessels. No evidence of antibody-mediated rejection. No evidence of recurrent IgA nephropathy.**

**Allograft kidney biopsy (6.5 months post-TPL):**

Light microscopy: Cortex with 7 glomeruli, two with fibrosis of the capsule. Focal interstitial fibrosis and tubular atrophy with lymphocytic and plasma cell infiltrates. Focal edema and infiltration of lymphocytes, macrophages and many plasma cells. Tubulitis with lymphocytes and plasma cells in tubules with intranuclear inclusions in tubular epithelium typical for polyomavirus. Two arteries with segmental sclerosis of the wall. Mild arteriolosclerosis.

Immunofluorescence (as stated in the original biopsy report): IgA, IgG, IgM and C3 negative in glomeruli. C4d negative in peritubular capillaries.

Immunohistochemistry (as stated in the original biopsy report): SV40 focally positive in nuclei of tubular epithelial cells.

Immunohistochemistry (performed for this study): HEV ORF2: Negative. SV40: Negative (despite morphology suggestive of polyomavirus infection and initial positive results outside).

Electron microscopy was not performed.

**Diagnosis: Polyomavirus nephropathy. Mild arteriolosclerosis. No evidence of antibody-mediated rejection. No evidence of recurrent IgA nephropathy.**

**Allograft kidney biopsy (4 months prior to death, Figure 1c):**

Light microscopy revealed 2 cores of cortex and medulla containing up to 8 glomeruli, two of which were hyalinized. Two glomeruli showed segmental sclerosis, one of those also endocapillary hypercellularity and prominent podocytes. Two glomeruli had mild mesangial expansion with minimal hypercellularity and mild increase in endocapillary mononuclear cells and neutrophils. Trichrome stain revealed diffuse, chunky mesangial and few glomerular basement membrane deposits, some suspicious for subepithelial “humps”.

The glomerular basement membrane showed rare holes and very rare splitting. Further findings included minimal interstitial fibrosis and tubular atrophy comprising <5% of the cortex, focal plasma cell-rich interstitial infiltrate involving less than 25% of the unscarred cortical parenchyma and moderate tubulitis with up to 10 leukocytes per tubular cross section, moderate arteriolar hyalinosis, and arteries with mild fibrointimal thickening. No peritubular capillaritis was found.

Immunofluorescence (as stated in the original biopsy report) revealed moderate (2+) granular mesangial and glomerular basement membrane positivity of IgG and C3. IgA, IgM, kappa and lambda were negative. C4d negative in peritubular capillaries.

Immunohistochemistry (performed for this study): Granular mesangial and glomerular basement membrane positivity for IgG and C3 (1+). IgA and IgM were negative. SV40: Negative.

HEV ORF2: Mild granular positivity in mesangium and glomerular basement membrane (1+).

Electron microscopy showed expanded mesangium with increased matrix and electron dense deposits. The lamina densa was irregularly thickened. There were subepithelial electron dense deposits, some of them hump-like, intramembranous and subendothelial deposits. Deposits were inhomogenously osmiophilic, but without substructure. There were no deposits in the tubular basement membranes.

**Primary diagnosis: Proliferative and focal sclerosing immune complex-mediated glomerulonephritis. Focal interstitial infiltrates and tubulitis, consistent with accompanying inflammation. No evidence of IgA nephropathy.**

**Diagnosis (upon re-evaluation and additional staining): Proliferative and focal sclerosing immune complex-mediated glomerulonephritis with positivity for HEV ORF2, consistent with HEV-associated glomerulonephritis. No evidence of IgA nephropathy, antibody-mediated rejection or polyomavirus nephropathy.**

**Allograft kidney biopsy (3 months prior to death, Figure 1c)**

Light microscopy revealed a core of renal cortex containing up to 9 glomeruli, 4 of which were hyalinized, 2 showed segmental sclerosis. The mesangium was mildly expanded, more than in the previous biopsy, with minimal, focal and segmental hypercellularity. There was very mild focal and segmental endocapillary hypercellularity with mononuclear cells and neutrophils. Trichrome stain revealed chunky mesangial and more glomerular basement membrane deposits, some suspicious for subepithelial "humps". The glomerular basement membrane showed rare holes and rare splitting. Minimal interstitial fibrosis and tubular atrophy comprising <5% of the cortex were found, focal severe arteriolar hyalinosis and arteries with mild fibrointimal thickening, but no peritubular capillaritis.

Immunofluorescence (as stated in the original biopsy report) revealed mild glomerular peripheral basement membrane and mesangial positivity for IgG (1+). IgM was negative.

Slides stained for IgA and C3 did not show any glomeruli.

Kappa and lambda light chains were positive.

Immunohistochemistry (performed for this study): Granular mesangial and glomerular basement membrane positivity for IgG (1+ to 2+) and C3 (1+). IgA was negative. IgM was not available. C4d was negative in peritubular capillaries.

HEV ORF2: Moderate granular to chunky positivity in mesangium and glomerular basement membrane (2+).

Electron microscopy was similar to the findings in the previous biopsy, but revealed even more deposits. Reticular aggregates were found in the cytoplasm of one endothelial cell.

**Primary diagnosis: Proliferative and focal sclerosing immune complex-mediated glomerulonephritis suggestive for viral infections.**

**Diagnosis (upon re-evaluation and additional stainings): Proliferative and focal sclerosing immune complex-mediated glomerulonephritis with positivity for HEV ORF2, consistent with HEV-associated glomerulonephritis. No evidence of IgA nephropathy, rejection or polyomavirus nephropathy.**

**Autopsy material of allograft kidney (11 years and 7 months after kidney transplantation, Figure 1c)**

Light microscopy showed more mesangial expansion and mild hypercellularity, still in a focal and segmental pattern. There was more endocapillary proliferation with mononuclear cells and some neutrophils, more splitting of the glomerular basement membranes. Trichrome stain revealed irregular deposits in the mesangium and glomerular basement membrane, some suspicious for subepithelial “humps”. Some glomeruli showed segmental sclerosis, some were hyalinized. There was focal interstitial fibrosis and tubular atrophy comprising approximately 20% of the cortex, moderate arteriolar hyalinosis not involving smooth muscle cells and moderate fibrointimal thickening in arteries, but no peritubular capillaritis.

Immunofluorescence (performed for this study) showed moderate to strong (2-3+) mesangial and glomerular basement membrane deposition of IgG, moderate (2+) mesangial and glomerular basement membrane deposition of C3, weak to moderate (1-2+) mesangial and glomerular basement membrane deposition of kappa light chains, moderate (2+) mesangial and glomerular basement membrane deposition of lambda light chains. Traces of IgM were detected in the mesangium and glomerular basement membrane. IgA was negative.

C4d was negative in peritubular capillaries.

For immunofluorescent double staining IgG/HEV ORF2, please refer to main manuscript, results section.

Immunohistochemistry (performed for this study) revealed moderate (2+) mesangial and glomerular basement membrane deposits of IgG and C3. IgA and IgM were negative. SV40 was negative.

HEV ORF2: Strong granular to chunky positivity in mesangium and glomerular basement membrane (3+).

Electron microscopy (performed for this study) was similar to the findings in the biopsies.

**Diagnosis: Proliferative and focal sclerosing immune complex-mediated glomerulonephritis with a membranoproliferative pattern and positivity for HEV ORF2, consistent with HEV-associated membranoproliferative glomerulonephritis with immune complexes. No evidence of IgA nephropathy. No evidence of rejection. No evidence of polyomavirus nephropathy.**

#### Patient 1 – liver specimen

**Autopsy liver (shown in Figure 1b):**

Light microscopic examination showed cirrhotic liver parenchyma with mild chronic active hepatitis, severe predominantly canalicular bile stasis and advanced autolytic changes.

Immunohistochemistry revealed patchy areas of hepatocytes positive for HEV ORF2 protein, mostly showing a cytoplasmic, but also a nuclear staining pattern.[1]

**Diagnostic: Chronic hepatitis E**

#### Patients 2-4 – kidney specimens

#### Patient 2

**Autopsy kidney (shown in Table 1):**

Histologic findings in kidney tissue obtained from patient 2 showed glomeruli with minimal endocapillary hypercellularity with mononuclear cells and pigmented tubular cast. Immunohistochemistry revealed moderate mesangial positivity for HEV ORF2 (2+), IgG (1+ to 2+), IgM (2+) and trace IgA and C3. Collectively, these findings were consistent with hepatitis E-associated proliferative immune complex GN and bile cast nephropathy. For immunofluorescent double staining IgG/HEV ORF2, please refer to main manuscript, results section.

#### Patient 3

##### Autopsy kidney (shown in Table 1):

Histologic findings in kidney tissue obtained from patient 3 showed autolytic changes with rare preserved cells in glomeruli and no preserved cells in tubules. The mesangium was expanded, consistent with diabetic glomerulosclerosis. Immunohistochemistry revealed moderate mesangial positivity for HEV ORF2 (2+), mild for IgG (1+), moderate IgM (2+), trace IgA and no C3. Collectively, these findings were consistent with hepatitis E-associated immune complex deposits without overt GN. In addition, some pigmented tubular casts were found, consistent with bile cast nephropathy. For immunofluorescent double staining IgG/HEV ORF2, please refer to main manuscript, results section.

#### Patient 4

##### Autopsy kidney (shown in Table 1):

Histologic findings in kidney tissue obtained from patient 4 showed rare preserved cells in glomeruli and no preserved cells in tubules. Inconspicuous glomeruli with autolytic changes. Immunohistochemistry revealed mild mesangial positivity for HEV ORF2 (1+) and trace IgM. IgG, IgA and C3 were negative suspicious for very mild hepatitis E-associated deposits without overt GN. In addition, some pigmented tubular casts were found, consistent with bile cast nephropathy. Material not suitable for EM.

#### Patients 2-4 – liver specimens

Histologic findings obtained in livers of patients 2 – 4 were recently described in Lenggenhager et al.[6], with patient 2 of this study corresponding to patient 35, patient 3 of this study corresponding to patient 27, and patient 4 of this study corresponding to patient 31.

### DETAILED DESCRIPTION OF THE METHODS

#### Histopathologic evaluation of tissue samples

Formalin-fixed, paraffin-embedded (FFPE) liver and kidney specimens were processed according to standard histologic methods including hematoxylin & eosin stain (H&E), liver specimens additionally with periodic acid-Schiff after diastase digestion stain (PAS-D) as well as a connective tissue stain such as Masson trichrome, kidney specimens additionally with periodic acid-Schiff (PAS), silver methenamine stain, elastic van Gieson-stain (EVG) and acid fuchsin orange G stain (AFOG).

Histologic slides including archived slides from prior kidney biopsies, liver autopsy specimens and newly stained slides were evaluated independently by experienced renal (BH, AG, HH, HY) or liver (DL, AW) pathologists, respectively.

#### Immunohistochemistry (IHC)

Distinct mouse monoclonal antibodies (mAbs) were used to target the different ORF2 protein isoforms: 1E6[1] and P3H2[7] for glycosylated (ORF2g/c) and nonglycosylated (ORF2i) isoforms, P1H1, P2H1 and P2H2 for HEV ORF2i only.[7, 8] FFPE cytospin material from Hep293TT cells replicating a full-length HEV genome HEV or the green fluorescent protein were used as positive and negative controls.[1] Visualization of the HEV ORF2 capsid protein was achieved using 1E6 (as described)[1] and P1H1, P2H1, P2H2 or P3H2 antibodies at a 1/250 dilution and included a 90 min pretreatment at 100°C with buffer CC1 and direct detection with OptiView Kit (Ventana).

For further immunohistochemistry on FFPE sections, standard procedures at the Department of Pathology and Molecular Pathology were applied using the Ventana BenchMark automated staining system with IgG, IgA, IgM, and C3 antibodies (Dako, Glostrup, Denmark).

#### Immunofluorescence

For IgG/HEV ORF2 double staining, we used either fresh frozen (6 µm thick) or FFPE (2 µm thick) tissue sections from kidney tissue. Sections were mounted on glass slides (SuperFrost Gold) and dried for 30 min at 37°C.

Fresh frozen sections were fixed in chemically pure acetone for 10 min. Slides were then air dried, pretreated for 5 minutes with a solution of 0.1% polyoxyethylene (20) sorbitan monolaurate (Tween 20) in tris-buffered-saline (TBS) and washed with distilled $H_2O$. FFPE tissue section were deparaffinized and pretreated with Tris/EDTA/Borat Buffer pH 9.0 for 30 min at 100°C.

Automatic staining was performed on a Leica Bond RX platform. Mouse monoclonal antibody clone 1E6 (Millipore Corporation, MAB8002) against the HEV ORF2 protein was incubated for 1h at a dilution of 1:125 followed by a mix of Alexa Fluor 546-conjugated goat anti-mouse antibody (Invitrogen BV, A11018) and FITC-conjugated Rabbit anti-Human IgG (Gamma chain, Diagnostic Biosystem, F008) for 1h at a dilution of 1:50. Following automated staining, the slides were hand washed in distilled $H_2O$. Tissue was covered with Vectashield® Antifade Mounting Medium with DAPI (Vector Laboratories, H-1200), covered with a coverslip and stored at 4°C until evaluation.

For single immunofluorescence on fresh frozen sections, standard procedures at the Department of Pathology and Molecular Pathology were applied using the Leica Bond automated staining system with IgG, IgA, kappa and lambda, C3, C4d antibodies (F008, F007, F001 and F002, F003, BI-RC4D respectively, Diagnostic Biosystem).

#### Transmission electron microscopy (EM)

Archived images from electron microscopy performed on the allograft kidney biopsies of patient 1, taken 11 years and 3 months (biopsy 3) as well as 11 years and 4 months post-transplantation (post-TPL) (biopsy 5) were re-evaluated. Additional ultra-thin sections from the archived grids were cut, stained and analyzed in a HITACHI TEM type H-7650 electron microscope.

Tissue samples of the allograft kidney from patient 1 were collected at autopsy and were fixed in 2.5% buffered glutaraldehyde for at least 12 h, post-fixed for 2 h in 1% osmium tetroxide, dehydrated in a series of graded ethanol solutions and propylene oxide and embedded in epoxy resin 48 h at 60° C. For comparison purposes, HEV negative tissue samples from the biopsy 1 (1 month post-TPL) of patient 1 were sourced from archive FFPE material.

Semi-thin sections (1 µm) were stained with methylene blue – azure. Ultra-thin sections (90 nm) were stained with UranyLess and lead citrate. Sections were examined in a HITACHI TEM type H-7650 electron microscope.

#### In situ hybridization (ISH) for HEV RNA

In situ hybridization for HEV RNA was performed as described using a commercially available probe designated V-HEV, targeting nucleotides 19-7257 of HEV-1 to -4 (#468111, Advanced Cell Diagnostics).[1]

#### Laser capture microscopy (LCM)

To perform mass spectrometry (MS) analysis of glomerular extracts from patient 1, glomeruli stained for HEV ORF2 protein (mAb 1E6) were dissected using the ArcturusXT™ LCM System (Thermo Scientific). Protein extracts from non-HEV-infected human kidney were used as controls.

Serial cuts of the formalin-fixed paraffin-embedded kidney specimen from patient 1 were prepared for LCM: one 2-µm thick section stained using mAb 1E6, two 10-µm thick sections stained using Cresyl violet placed on polyethylene naphthalate (PEN) membrane glass slide and kept overnight in 4°C.[9, 10] Areas of interest were identified by overlapping the HEV ORF2+ glomeruli with the matching structures on the 10-µm thick section on PEN membrane glass slides. Isolation of glomeruli was verified by microscopic examination of the LCM cap as well as the excised region (Figure 2c). Two caps containing 40-50 glomeruli each were collected per case. After excision, the caps containing tissue were transferred to a 1.5-mL centrifuge tube (Eppendorf ®Safe-Lock tubes) and frozen at −20°C until further processing.

#### Laser-captured sample preparation for mass spectrometry (MS)-based protein identification

For protein extraction, sterile blades and forceps were used to peel off the thermoplastic membranes containing laser-captured glomeruli from the LCM cap, which were then transferred into a sterile Eppendorf® Safe-Lock tube.

Samples were submitted to the Functional Genomic Center Zurich for mass spectrometry analysis. Briefly the samples were lysed by adding 4% SDS lysis buffer and incubated for 60 min at 95°C, 1 min HIFU (High Intensity Focused Ultrasound), 10 min sonication, 30 min of additional heating at 95°C, HIFU, sonication and max spin for 5 min. The samples were digested using trypsin solution on a thermo shaker at 37°C overnight. The digested samples were dried and dissolved in 3% aqueous Acetronitrile and 0.1% formic acid before being transferred in vials for liquid chromatography-mass spectrometry analysis.

The acquired MS data were processed for identification using the Maxquant search engine. The spectra were searched against the HEV ORF2 protein sequence merged with the Homo sapiens protein database Swissprot, and also against interstitial markers (cytokeratin 7, complement 3, and collagen 1A1) and specific glomerular marker podocin.[2, 3] Results were summarized in the Proteome Software Scaffold under very stringent settings (1% protein false discovery rate [FDR], a minimum of 2 peptides per protein, 0.1% peptide FDR*)* and expressed as averaged total spectrum counts ± s.d. (Supplementary Table 2).

#### Image processing

Slides with histologic / histochemical stains, immunohistochemistry and ISH were evaluated by light microscopy and scanned with a resolution of 0.250 µm per pixel (on a Hamamatsu Nanozoomer 2.0 HT whole slide imager) for evaluation by digital microscopy (NDP.view2 viewer software, Hamamatsu Photonics K.K.). Regions of interest were exported as tif. files from scanned slides, from images acquired from a Zeiss Axio Scope.A1 equipped with a Zeiss Axiocam 100 color camera controlled via ZEN 2.3 lite Software, or from images acquired with an Olympus BX43 microscope equipped with an Olympus DP-43 digital camera controlled via Olympus cellSens software.

Immunofluorescence images were acquired with an upright fluorescence microscope (AxioImager.Z2 controlled by ZEN Blue software; 89 North Photofluor LM-75 light source, and Axiocam 503 mono camera; Zeiss, Jena, Germany), equipped with the following objectives: 20x (NA 0.5, Plan-NEOFLUAR), 40x (NA 1.4 oil, Plan-APOCHROMAT), and 100x (NA 1.45 oil, Plan-APOCHROMAT) objectives. This setup provides an excellent spatial resolution (nominally about 200-nm lateral resolution in our study; pixel size was 45.4 nm for 100x objective) comparable to confocal microscopy.[11] High resolution images were taken with the 100x objective using the ApoTome.2 module with deconvolution (grid 5 lp/mm; section thickness 0.7 µm). We used Vysis Abbott Chroma filter sets (Blue: excitation (ex) 335-383 nm, emission (em) 420-470; green: ex 481-507 nm; em 521-551 nm; red: ex 534-556 nm, em 574-606 nm). Control experiments with separate single-channel IgG or HEV ORF staining revealed negligible cross-talk between green and red fluorescence channels.

Co-localization of IgG and HEV ORF was quantified using Fiji[12] and the JACoP ImageJ plug-in.[13] For analysis of entire glomeruli, images taken with the 20x objective were cropped to a rectangle around the glomeruli's outlines. For the 100x images, subareas within individual glomeruli of 45-85 µm side lengths were selected. Pearson's correlation coefficient (PCC) was calculated using Costes approach for automatic thresholding.[14, 15] To test for significance of PCCs we used Costes randomization (1000 rounds) on each image pair with block sizes of 4 pixel (0.908 µm) for 20x and 10 pixel (0.454 µm) for 100x. P-values were all highly significant ($p < 10E-10$). We also calculated the Manders coefficients M1 and M2[16], which for patient 1 were between 0.64 and 0.98 for 20x images ($M1 = 0.84 \pm 0.08$, $M2 = 0.83 \pm 0.09$; mean ± s.d., $n = 25$ glomeruli) and between 0.22 and 0.83 for 100x images ($M1 = 0.58 \pm 0.14$, $M2 = 0.72 \pm 0.12$; mean ± s.d.), consistent with high degrees of co-localization. Thresholds for M1 and M2 analysis were set manually.

For ultrastructural evaluation, ultrathin sections were examined using a transmission electron microscope (HITACHI H-7650). Regions of interest were exported in tif format. Figures were created using Adobe Photoshop software.

#### Western blotting

Autopsy kidney and liver tissue samples from patient 1 and normal kidney tissue sample from non-HEV-infected patient were homogenized in RIPA buffer (R0278-50ML, Sigma-Aldrich,) containing phosphatase inhibitor and protease inhibitor cocktail tablets. Protein extracts were separated by SDS-PAGE on a 4-20% precast polyacrylamide gel (#4561094, BioRad), transferred to a PVDF membrane according to the manufacturer's protocol and analyzed by Western blotting. The antibodies used were directed against HEV ORF2 protein (mouse monoclonal antibody 1E6, MAB8002, Millipore), PAX8 (10336-1-AP, Proteintech), Arginase-1 (SP156, Cell Marque), and β-Actin (4970T/S, Cell Signaling). Protein expression was detected by chemiluminescence and gel-imaging system (Bio Rad).

#### Molecular testing: qRT-PCR for HEV

For HEV RNA detection, a qRT-PCR protocol was applied to fresh frozen and FFPE tissue specimens. To roughly determine RNA quality and quantity, qRT-PCR for the housekeeping gene β-actin (ACTB) was performed.

RNA was extracted from three 10 µm thick tissue sections from each organ using either the RNeasy FFPE Kit (Qiagen) or RNeasy Plus Mini Kit (Qiagen) depending on the sample type according to the manufacturer's instructions. One µg of mRNA was reverse transcribed using the iScript Reverse Transcription Supermix for RT-qPCR (Bio Rad) according to the manufacturer's instructions. Hep293TT cells replicating a full-length HEV genome[1] briefly fixed with 10% neutral buffered formalin and embedded in paraffin as cellblocks, were used as a positive control and treated in the same way as patient biopsies.

HEV qPCR reactions were performed in a final volume of 10 µl containing 40 ng cDNA, 200 nM forward primer/HEV probe/reverse primer mix, and TaqMan Fast Advanced Master Mix

(Thermo Fisher). Primer and probe sequences are defined in the table below. qPCR assays were performed in an Abi Viia7 qPCR 384 Detection System (Thermo Fisher) using the following program: 95°C for 20 sec, followed by 40 cycles of 95°C for 1 sec, 60°C for 20 sec. Three control reactions were run in parallel: (1) Positive control reactions contained 10-fold serial dilutions (C1: 4 ng, C2: 0.4 ng, C3: 0.04 ng) of the cDNA obtained from HEV-replicating Hep203TT cells. A Ct threshold value for the HEV amplicon was set at 0.05 $\Delta$Rn. HEV detection limit was set at the Ct value of the lowest dilution of the positive control, which was normally at the Ct 35. (2) As a control of the cDNA quality, all patient samples were also assayed with FAM-labelled $\beta$-actin (ACTB) TaqMan assay Hs01060665_g1 (Thermo Fisher). A Ct threshold value for the ACTB amplicon was set at 0.1 $\Delta$Rn. A sample passed the quality control, if the Ct value of the ACTB amplicon was less than 35. (3) Water (no template control) was run as one of the samples to exclude reagent contaminations.

Primer and probe sequences

| Name | Sequence 5' - 3' |
|---|---|
| JVRSHEV_F1 | CGGTGGTTTCTGGGGTGAC |
| JVRSHEV_F2 | GCAGTGGTTTCTGGGGTGAC |
| JVRSHEV_F3 | GCGGTGGTTTCTGGAGTGAC |
| JVRSHEV_F4 | GCGGTAGTTTCTGGGGTGAC |
| JVRSHEV_R1 | CAAAGGGGTTGGTTGGATGAA |
| JVRSHEV_R2 | GAAGGGGTTGGTTGGATGAA |
| JVRSHEV_R3 | CGAAAGGGTTGGTTGGATGAA |
| JVRSHEV_R4 | AAGGGGTTGGCTGGATGAA |
| JVRSHEV_R5 | CGAAGGGATTGGTTGGATGAA |
| JVRSHEV_Pt | (FAM)-ATTCTCAGCCCYTYGC-(BHQ) |

A sample was considered as HEV positive or negative if the average Ct value of the HEV amplicon was below or above the average Ct value of the C3 positive control, respectively.

#### Molecular testing: HEV genotyping

Plasma sample was subjected to HEV genotyping following amplification of HEV ORF2 (592 bp) sequence, as recently described.[17] Genotype and subtype assignment were performed using the HEVnet genotyping tool (https://www.rivm.nl/mpf/typingtool/hev/).

#### Serological assay for anti-HEV IgG and IgM

Serum sample obtained in Cantonal Hospital Aarau was subjected to VIDAS® ANTI-HEV IgG and IgM assay (BioMérieux, France). Sample from University Zürich Hospital was assessed using Enzyme ImmunoAssays for the determination of IgG and IgM antibodies to hepatitis E Virus provided by Dia. Pro Diagnostic Bioprobes Srl (Italy).
